# The return to water in ancestral *Xenopus* was accompanied by a novel mechanism for producing and shaping vocal signals

**DOI:** 10.1101/360255

**Authors:** Ursula Kwong-Brown, Martha L. Tobias, Damian O. Elias, Ian C. Hall, Coen P.H. Elemans, Darcy B. Kelley

## Abstract

Species-specific vocal signals allow listeners to locate potential mates. During the tetrapod transition from water to land, lungs replaced gills, allowing expiration to drive sound production. Several groups, *e.g*. cetaceans and some frogs, then returned to water. Here we explore how air-driven sound production changed upon re-entry and how essential acoustic information on species identity was preserved in the secondarily aquatic frog *Xenopus*. We filmed movements of cartilage and muscles during evoked sound production in isolated larynges. Our results refute the current theory for *Xenopus* vocalization, cavitation, and instead favor sound production by mechanical excitation of laryngeal resonance modes following rapid separation of laryngeal arytenoid discs. The resulting frequency resonance modes (dyads) are intrinsic to the larynx rather than due to neuromuscular control. We show that dyads are a distinctive acoustic signature across species. While dyad component frequencies overlap across species, their ratio is shared within each *Xenopus* clade and thus provide information on species identity, potentially facilitating both conspecific localization and ancient species divergence.

## Introduction

In the transition from water to land in early tetrapods, lungs replaced gills for respiration.^1^ Many current tetrapods use air movement to empower specialized vocal organs such as the larynx of frogs and mammals and the syrinx of birds.^2^ The resulting sounds are shaped by a combination of vibrating elements and cavity resonances to voice different acoustic qualities that convey sex, age, species, emotional state and even intent.^3^ While voice provides essential information for social interactions, we know surprisingly little about how vertebrate vocal organs create these complex acoustic features. In particular, when an ancestral tetrapod leaves the land and vocalizes underwater without air movement, how are communication sounds produced and then shaped to maintain essential social information, and how do they diversify during speciation? Frogs in the secondarily aquatic genus *Xenopus*^4^ present an informative system for addressing these questions.

In *Xenopus*, social communication is dominated by vocal signaling.^5^ Males in each species produce distinctive advertisement calls underwater whose acoustic features inform species identity.^4^ These calls consist of a series of sound pulses that form species-typical temporal patterns and characteristically include two dominant frequencies (DFs).^3,6^ The sound pulses that comprise *Xenopus* calls are produced in the larynx,^7^ a vocal organ interposed between the nasal and buccal cavities and the lungs (Fig. 1A). Vocal folds are absent^8^ and the glottis gates air flow to and from the lungs.^9^ The larynx consists of a cricoid frame or “box” of hyaline cartilage flanked bilaterally by bipennate muscles. These insert anteriorly, via a tendon, onto paired, closely apposed arytenoid cartilage discs (Fig. 1B) whose medial faces are coated by mucopolysaccharide secreted by adjacent cells.^10^ The discs are suspended in elastic tissue composed of elastic cartilage and elastin fibrils.^10^ Electrical stimulation of laryngeal muscles or nerves results in species specific sound pulses, both *in situ* and *ex vivo*.^10,7^ Sound is thus produced without air flow or vocal folds. In *X. borealis*, while separations of the paired arytenoid discs accompany sound pulses,^10^ how disc motion results in sound production has not been resolved.

**Fig. 1.**
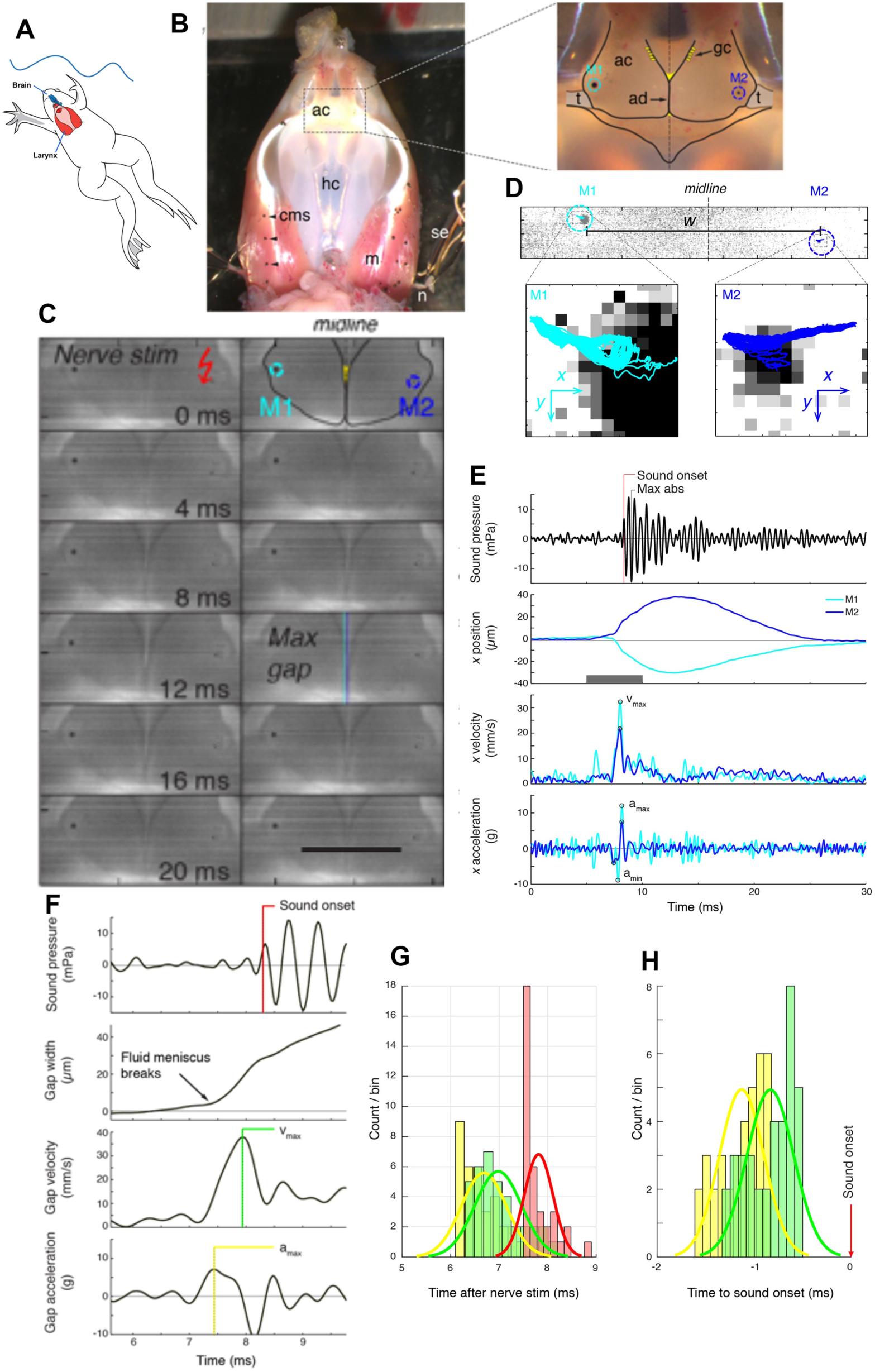
A) Xenopus call while submerged. B) Ventral aspect of the isolated *Xenopus laevis* larynx, a cricoid box of hyaline cartilage (hc) flanked by muscles (m). Sounds are produced by the separation of apposed arytenoid cartilage (ac) discs in response to muscle (m) contraction evoked by bilateral nerve (n) stimulation via suction electrodes (se). Carbon microspheres (cms) placed on the surface track muscle and cartilage positions. Panel at right: a spherical carbon bead (M1, M2) is placed atop each disk. Goblet cells (gc) secrete mucopolysaccharide onto the disc surfaces. Muscles insert via a tendon (t) onto the posterolateral edge of each arytenoid disc (ad). C) Still photo of arytenoid motion filmed at 10,000 fps illustrated at 1 ms intervals. Nerve stimulation occurs in top left image. Scale bar 1mm. D) Upper panel: Higher magnification images of each bead in B) during filming at 44,000 fps. Lower panel: The small motion of each bead (cyan, left; blue, right) during 40 consecutive stimulations at 40 Hz. E) Sound (top panel, corrected for time of flight) and the position, velocity and acceleration of the two beads (color coded as in D) during a single pulse. Nerve stimulation occurs at t=0. F) Kinematic data for gap width (w) in C) in relation to sound onset. While the precise onset (red line) is hard to determine due to acoustic noise, sound production follows peak velocity (green) or acceleration (yellow). G) The timing of sound onset (red), gap peak acceleration (yellow) and peak velocity (green) during 40 consecutive clicks for one larynx relative to nerve stimulation. H) Sound onset relative to gap peak acceleration (yellow) and peak velocity (green) during 40 consecutive clicks for the larynx in G). I) Sound production follows peak acceleration and velocity after 0.85±0.33 and 0.51±0.33 ms respectively for three individual larynges.

An unusual mechanism – implosion of air bubbles or cavitation^10^ - is the currently accepted^11,12^ hypothesis for underwater laryngeal sound production in *Xenopus*. In this scenario, the high velocity separation of the arytenoid discs causes formation of bubbles that then implode and produce sounds. Cavitation bubbles are known to produce hydrodynamic propeller noise^13^ and the “snaps” of some species of shrimp.^14^ However, *a priori* cavitation - creating “a bubble between the discs at a pressure below ambient … that … implodes as air rushes into the cleft at high speeds, producing a click” ^10^ - seems an unlikely cause of sound production in *Xenopus*. The small film of fluid between the arytenoid discs should allow neither high velocity flow nor bubble formation, and bubbles have not yet been observed. Additionally, cavitation bubble collapse produces a high amplitude pressure pulse (~50-100 kPa), several orders of magnitude louder than radiated sound pressure of *Xenopus* sound pulses. Finally, the duration of pressure transients produced by collapse of cavitation bubbles are in the microsecond range, rather than the millisecond range of *Xenopus* sound pulses.

## Results

### Mechanism of sound pulse production

To empirically test the cavitation hypothesis, we filmed isolated *X. laevis* larynges during sound pulse production evoked by stimulation of the laryngeal nerve.^7^ As reported previously,^10^ disc movements accompanied sound production (Fig. 1C,E,F; Supplementary Video 1). To track the position of the arytenoid discs, we placed perfect carbon microspheres over the discs (Fig. 1B,C) and computed disc position at subpixel resolution by interpolating the 2D intensity correlations of the spheres with each image (See **Methods**, Fig. 1D). This approach revealed disc position at speeds of up to 44,000 fps and allowed us to calculate disc velocity and acceleration profile in relation to sound pressure (Fig. 1E-H).

Nerve stimulation first produces isometric contraction of the bipennate muscles during which the arytenoid discs remain apposed. In favorable preparations, a fluid layer could then be observed retreating from the medial surface of the discs with increasing isometric force. This observation suggests that the discs are kept together by a capillary binding force through a liquid bridge. When bilaterally exerted muscle force overcomes the binding force, the liquid bridge ruptures (Fig. 1F) and the discs separate rapidly with mean gap peak acceleration of 13.5±9.1 *g* (N=3; range: 6.4-23.8 g, where 1 *g* = 9.82 m/s^5^) and mean peak velocity of 50.3±21.9 mm/s (N=3; range: 35.5-75.4 mm/s) (Fig 2C-D). Disc peak deceleration (10.6±5.2 g; N=3; range: 6.5-16.4 g) occurs only 0.57 ±0.10 ms after peak acceleration. In this short time, the gap between the discs enlarges to 18.6±5.5 μm (range: 14.1-24.8 μm), a value that is 27.5±8.0% (range: 22.8-35.6%) of the maximum gap width of 72.6±34.5 μm (N=3; range: 39.6-108.4 μm μm). Peak acceleration and velocity precede sound onsets by 0.85±0.33 and 0.51±0.33 ms respectively (Fig.1 E,F). The interval between disc separation and subsequent sound radiation reveals that these events are clearly associated. The delay (Fig 1F) between disc acceleration and velocity shows less variation (0.34±0.06 ms, N=3) indicating that sound onset timing after disc separation - rather than disc kinematics - is more variable between preparations.

**Fig. 2.**
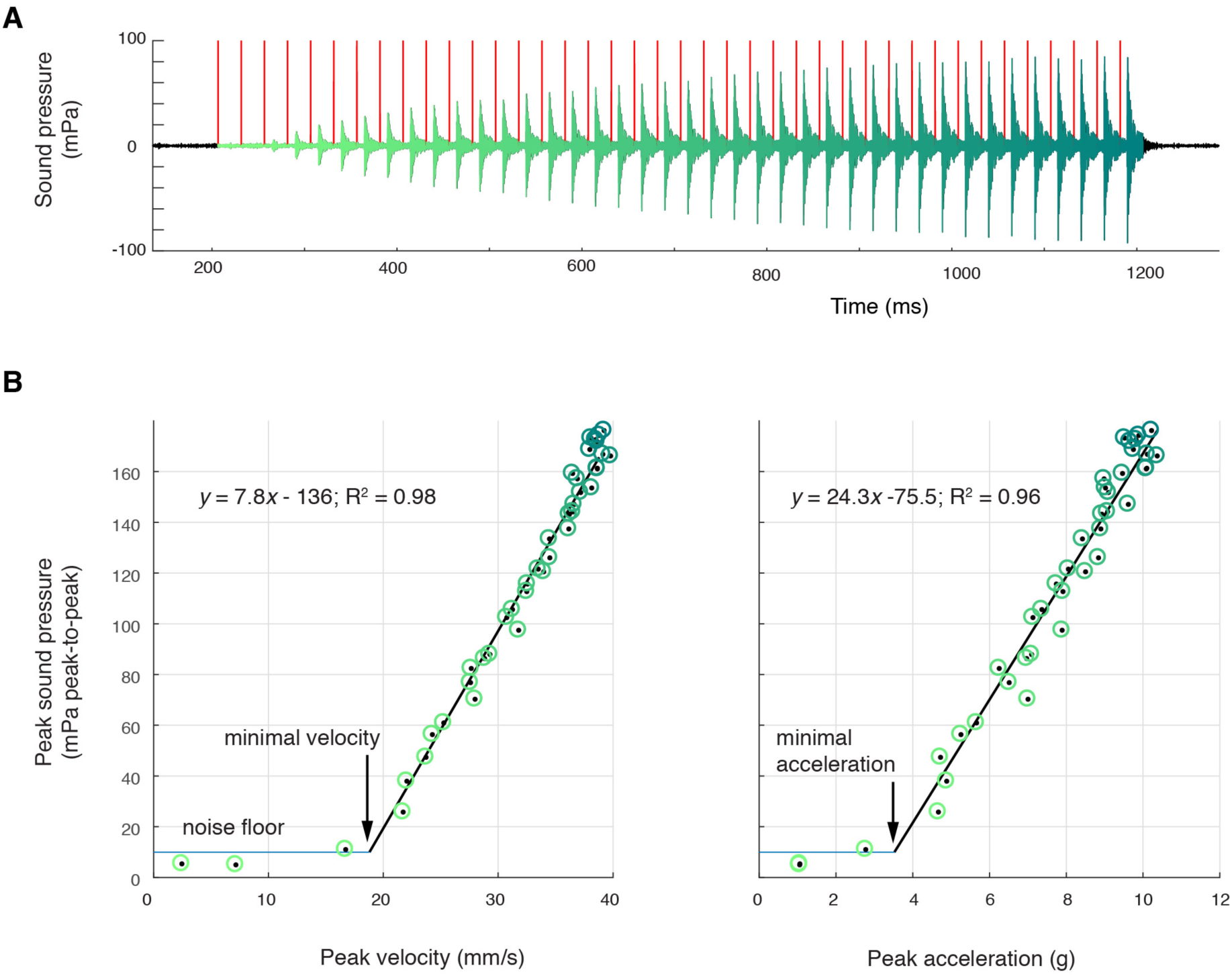
Sound amplitude correlates linearly with disc kinematics, A) Sound pressure across 40 consecutive stimulations (red vertical lines) at 40 Hz (green, first stimulus; blue, last stimulus). B) Peak sound pressure per stimulus (color-coded as inl A) increases linearly with peak velocity (linear regression for three individuals: y=5.2×-210.5, R2 = 0.96; y=1.8×-35.1, R2 = 0.89; y=7.8-136.3, R2 = 0.98) and peak acceleration (linear regression for three individuals: y=14.3×-106.9, R2 = 0.91; y=5.6×-6.1, R2 = 0.89; y=24.3-75.5, R2 = 0.96). A minimal velocity and acceleration is required before sound is radiated.

At 40Hz nerve stimulation rates (within natural call sound pulse rates^6^), the first few stimuli do not result in sound pulse production (Fig. 2A) corroborating previous results.^7,15^ Subsequently, over multiple stimulations, peak-to-peak sound pressure increases linearly with both peak velocity and acceleration and reaches a maximum received level of ~180 mPa ptp (at 44 mm) in all preparations (Fig 2B). Below a minimal peak velocity of 28±12 mm/s and minimal peak acceleration of 4.6±3.1 g, no sound is detected, suggesting a threshold disc velocity or acceleration required to produce sound. When water was introduced via the glottis into the supradisc space, though peak velocity reached this threshold, peak acceleration did not (Supplementary Figure Sl2), and no sound was produced, corroborating earlier observations in *X. borealis*^1^

These observations do not support the cavitation hypothesis because we do not observe bubble formation, sound pressures are six orders of magnitude too low (mPa vs kPa), and the onset of sound production is three orders of magnitude too slow (ms vs μs).

### Dyad ratios are shared within each clade

We have previously reported^7^ that sound pulses produced *ex vivo* in male *X. laevis* larynges include species-specific frequencies. To determine whether spectral features of calls reflect species-typical laryngeal features, we first recorded male advertisement calls from representative *Xenopus* species. In the three clades of this sub-genus - L (which includes *X. laevis)*, M (which includes *X. borealis*) and A (which includes *X. andrei* and *X. amieti*)^4^ - each repeated sound pulse in the male advertisement call includes a higher dominant frequency (DF2) and a lower frequency (DF1) band^6^, termed dyads (Fig. 3A, C, D). All but one species produce advertisement calls made up of sound pulses containing two, simultaneously produced dyads. The exception is *X. allofraseri* in which pulses contain harmonic stacks. There is a broad range for both the lower and the higher dominant frequency across *Xenopus* (Fig. 3C, D). Either DF1 or DF2 can be shared by different species. However, in contrast to individual frequencies, the ratio of DF2:DF1 is specific to, and highly conserved within, each clade (Fig. 3E). The mean ratio (see SupplementaryTable 1) for A clade species is 1.34 (±0.01), for L clade species is 1.24 (±0.03), and for M clade species is 2.02 (±0.01). Exceptions include *X. wittei* in the A clade (1.19, not 1.34) and *X. laevis* South Africa in the L clade (1.14, not 1.24). Thus, species in different clades can share either DF1 or DF2 but not the distinctive ratio.

**Fig. 3.**
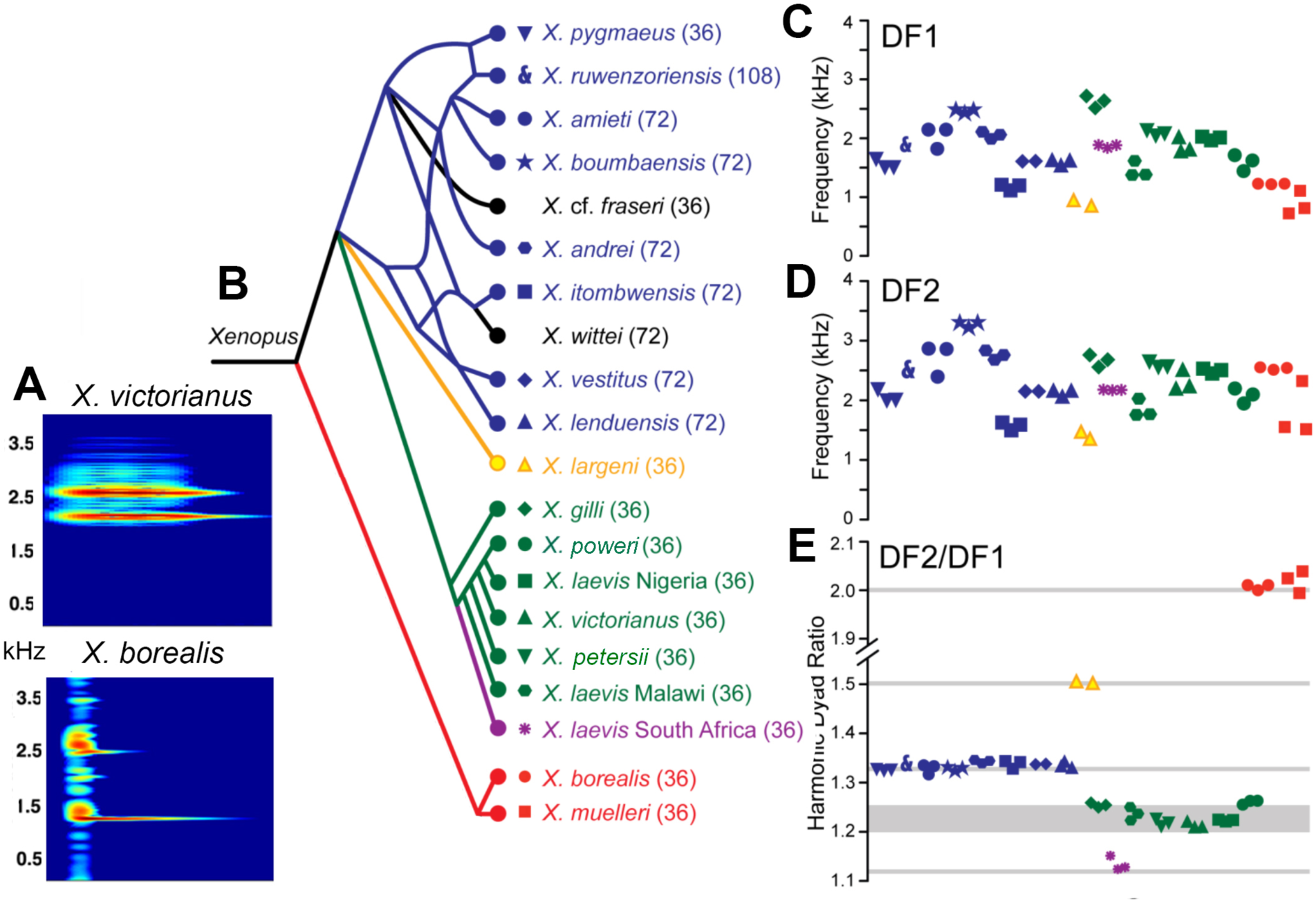
Harmonic dyad ratios are specific to, and highly conserved within, each *Xenopus* clade (blue: Clade A, green: Clade L; red: Clade M). **A**) Each sound pulse in an advertisement call includes two dominant frequencies (DFs). Sound spectrograms for three individual sound pulses in advertisement calls of *X. victorianus* (L clade); duration of each pulse is ~30ms. Sound spectrogram of one sound pulse in *X. borealis* (M clade) advertisement call, ~ 50ms duration. **B**) Phylogenetic relationships of *Xenopus* species in this study. Clades and the DF2:DF1 ratios are: 1.34 (blue, A clade), 1.21-1.26 (green, L clade), and 2.0 (red, M clade). *X. allofraseri* (black) sound pulses are harmonic stacks. *X. laevis* South Africa (purple) and *X. wittei* (black) sound pulses are exceptions to their species group ratios. The ploidy level (number of chromosomes) is in parentheses; the DF2:DF1 (dyad) ratio for individual male calls for each species is indicated by a unique combination of symbol and color (**B-E**). The value of the lower dominant frequency (DF1; **C**) and the higher dominant frequency (DF2; **D**) respectively, in advertisement call sound pulses across *Xenopus*. **E**) Harmonic dyad ratios fall into 3 major bands, one for each clade.

### Dyads are intrinsic to the larynx

To determine how dyads are produced, we first confirmed that our recordings of sound pulses were free of possible acoustic artifacts produced by interactions with the recording tank. Using a hydrophone and laser Doppler vibrometry, we obtained simultaneous recordings of sounds and body vibrations from a calling male under the same conditions in which we obtained the data in Fig. 3. The same dyad was present in both sounds and vibrations from a vocalizing male (Fig. 4A). Thus, the dyads reported here are produced by the frog. To determine whether dyads are produced solely by the larynx (without contributions from other organs), we recorded sound and vibrations produced by isolated larynges in response to nerve stimulation (Fig. 4B). Isolated larynges produced the same dyads as the intact animal. Thus, the generation of dyads is intrinsic to the larynx and is not influenced by extra-laryngeal tissues.

**Fig. 4.**
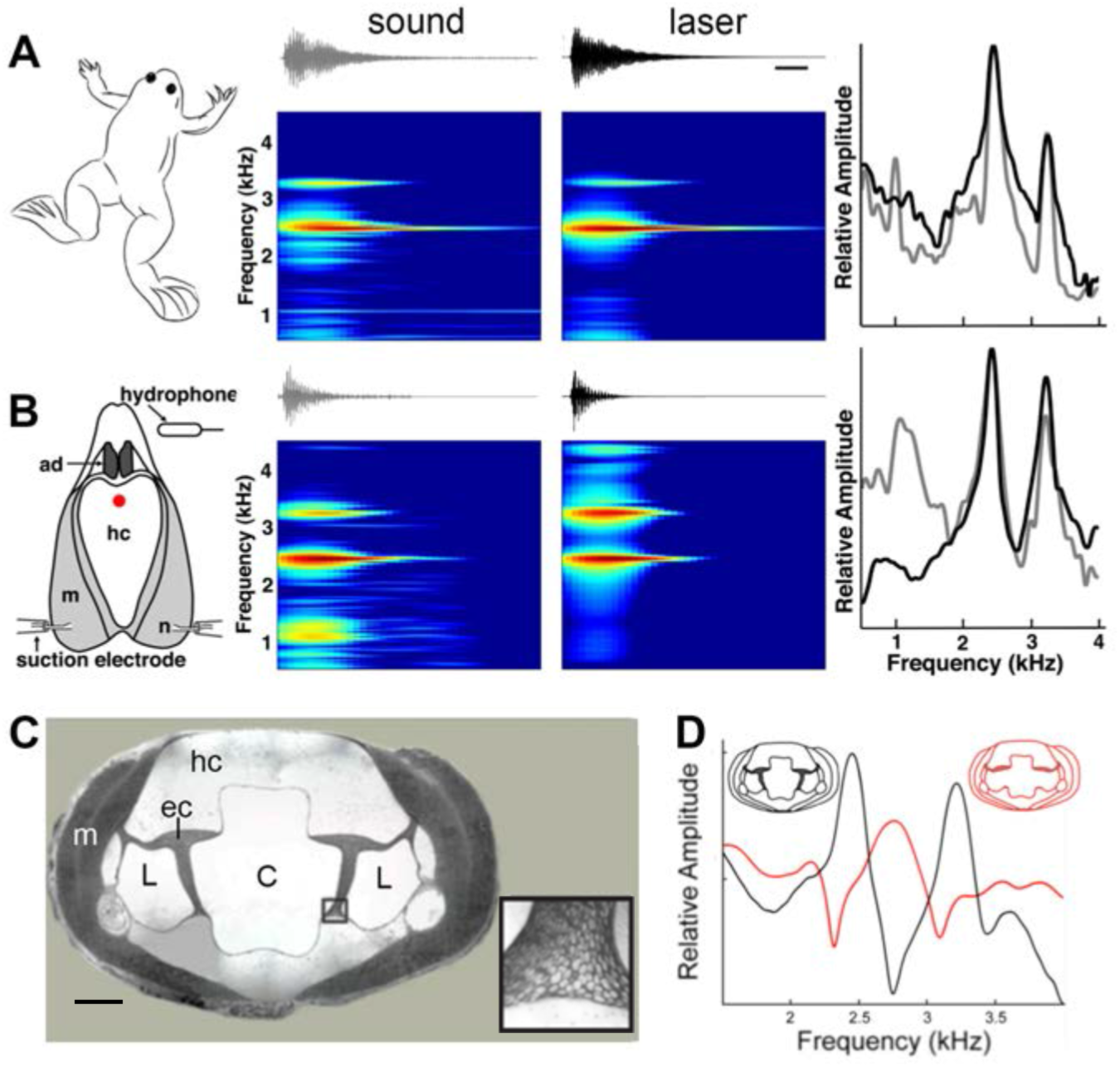
A singing male and an isolated larynx with intact elastic cartilage septa produce the same dyad in sound and laser recordings **A**) Recordings from a *X. boumbaensis* male. Spectrograph (frequency vs time) of one pulse obtained from sound and laser vibrometry recordings. Red spot indicates the location of the laser recording. Scale bar: 12ms. At right: Power spectra from the sound (grey) and laser (black) spectrographs. **B**) Recordings from the isolated larynx of a *X. boumbaensis* male. Spectrograph (frequency vs time) of one pulse obtained from sound and laser vibrometry recordings, At right: Power spectra from the sound (grey) and laser (black) spectrographs. Note that the broad band frequency peak ~1 kHz in the sound recordings (gray) is not present in the laser recordings (black) and is thus an artifact of recording conditions (glass aquaria or Petri dish). Spectrographs are color graded to show increased intensity (blue to red). **C - D** Intact elastic cartilage is required for the production of frequency dyads. **C**) Transverse section of an osmicated, epon-embedded male *X. laevis* larynx just anterior to the nerve (n) entry point in **B**; dorsal is up. The hyaline cartilage (hc) frame is flanked laterally by paired, bipennate laryngeal dilator muscles (m). Elastic cartilage sheets divide the lumen of the larynx into a central chamber (C) and two symmetrical lateral chambers (L). Elastic cartilage is recognizable by its unique, lace-like cellular morphology (inset). Scale bar 1mm. **D**) When elastic cartilage of the isolated larynx is disrupted bilaterally (red) via puncture (see Supplementary Figure SI1), the two DFs characteristic of the intact larynx (in black) are replaced by a single, intermediate DF in red).

The ability of tissues containing elastic cartilage - such as the pinna - to deform and reform rapidly suggested that this tissue might be essential to producing dyads. The interior of the larynx (Fig. 4C) includes a central, air-filled chamber (C) separated from smaller, lateral chambers (L) by elastic cartilage septa (Fig. 4C, inset). We isolated the larynx and drilled a small hole into the middle of the dorsal hyaline cartilage through which a pin was used to puncture the elastic cartilage on both sides. After puncture, the elastic cartilage was no longer structurally intact (Supplementary Fig. Sl2C). In the three species examined - *X. boumbaensis* (A clade; n = 6), *X. victorianus* (L clade, n = 3) and *X. laevis* South Africa (L clade-DF2/DF1 exception; n = 6) - the hole in the hyaline cartilage by itself (Sl2A) did not affect the frequencies produced by motor nerve stimulation (Sl2A). However, puncturing the elastic cartilage either abolished the DF peaks by shifting the two narrowly tuned bands to a single, intermediate, broader band frequency (10/15 larynges; Fig. 4D), abolished one peak and broadened the other (2/15), abolished both peaks (2/15) or shifted peaks to a different ratio (1/15). In *X. borealis*, “opening” the cricoid box by removing a rectangular portion of the ventral laryngeal wall detunes the larynx.^10^ This detuning could also have been due to disruption of elastic cartilages. Because the elastic cartilage partitions the internal air chambers of the larynx, the puncture created a single air space. To examine a potential role for the internal air spaces, we reduced the volume of the lumen in isolated *X. laevis* male larynges by inserting a large bead or replaced air with helium (as in a previous study^10^). We also placed weights on ventral surface of the larynx. None of these manipulations affected DF1, DF2 or the DF2/DF1 ratio. These observations indicate that the dyads do not reflect the volume of the internal chambers or the mass of the cricoid cartilage.

## Discussion

Our data do not provide support for the prevailing^10–12^ cavitation hypothesis for sound production in *Xenopus*. Two alternative mechanisms that could account for the association between disc movement and sound production are: 1) acoustic excitation by a rapid pressure drop between the discs or 2) disc movement-associated mechanical excitation of the larynx. Sound pressure reduction between the discs may produce a propagating sound pressure wave, exciting air cavity resonances within the larynx. This mechanism should result in a bipole sound source with strong directional radiation pattern.^16^ Because of impedance mismatch between the air cavities and cartilages of the larynx, however, this mechanism would produce a poor, low intensity sound. In addition, replacing air with helium should alter the frequency distribution of cavity resonances,^17^ an effect not observed in this or a previous^10^ study.

Alternatively, separation of the arytenoid discs might mechanically excite vibration of laryngeal elements. This mechanism would result in a monopole sound source – the entire larynx - with a more omnidirectional radiation pattern, effectively coupled to the medium, producing a more intense sound. The sound pressure produced by such a vibrating monopole structure depends on its space and time averaged velocity,^18^ which is consistent with our observations of a linear correlation between disc velocity and sound pressure, and the minimal velocity required for sound production. This mechanism is also consistent with previous experiments in which splitting “the elastic sac surrounding the discs”^10^ prevented sound pulse production. We thus favor the second explanation and propose that disc movements specifically excite vibration of the elastic tissues surrounding the discs (Supplementary Fig. Sl3).

As key features of the *Xenopus* larynx - including lack of vocal folds and modification of the laryngeal box and cartilages - are shared across Pipid species, ^8,19^ this proposed mechanism of underwater sound production may also be shared. In *Xenopus*, water is prevented from entering the larynx during underwater calling by inhibition of glottal motor neurons,^20^ thus ensuring the attainment of the disc acceleration values required for sound production identified here. The sound-protection afforded would not be required in another pipid, *Hymenochirus merlini*, that has reverted to calling in air,^11^ presumably through an open glottis. We predict that *H. merlini* calling is also powered by disc separation rather than air flow.

Our experimental results furthermore support the hypothesis that arytenoid disc movements subsequently excite two natural vibratory resonance frequencies of the larynx itself, the harmonic dyads. These dyad resonance frequencies require intact elastic cartilage septa that separate the central laryngeal lumen from the lateral laryngeal chambers. Both species-specific individual DFs and the clade-specific dyad ratio are thus intrinsic to the larynx rather than the result of laryngeal or respiratory muscle modulation by neural circuitry. Which as yet unidentified characteristics of laryngeal tissue geometry and properties result in species-specific DFs and their ratios remain to be determined but are likely to reflect a common tuning mechanism in descendants of ancestral *Xenopus* species.^21,22^

Three mechanisms have been identified for producing and shaping vertebrate laryngeal vocalizations: the myoelastic aerodynamic theory (MEAD), active muscle contraction (ACM) and intralaryngeal aerodynamic whistles. The MEAD mechanism^23^ explains the physical basis for sounds produced by isolated larynges of mammals and terrestrial frogs as well as by syringes in birds.^2^ The ACM mechanism requires motor neuron-driven contraction of intrinsic laryngeal muscles to produce, for example, purring in cats.^24^ Intralaryngeal aerodynamic whistles produce ultrasound in mice and probably all murine rodents^25,26^ All of these mechanisms require mass flow to produce vocalizations and thus neither explains how *Xenopus* call. The return to water in ancestral *Xenopus* was instead accompanied by a novel mechanism for laryngeal sound production: disc-induced vibrations excite laryngeally intrinsic resonance modes - dyads - shaped spectrally by material properties of elastic cartilage septa. Thus, the evolutionary change that allowed sound to be produced underwater, without mass airflow, in Pipids is not only responsible for production of sound pulses but also their spectral features. Species-specific, rhythmic activity patterns of laryngeal motor neurons drive the precise temporal pattern of laryngeal muscle contractions responsible for the arytenoid disc separations that produce vocalizations.^27–29^

In frogs, as in other vocal vertebrates, acoustic features of male advertisement calls contain information on species identity.^30^ This is also true for *Xenopus*; other call types - such as the male release call - vary little between species.^31^ Information on species identity can serve to reduce interspecific mating and costs associated with hybrid offspring, including male sterility, that lead to restricted gene flow and speciation.^32^ Across *Xenopus*, temporal patterns of the advertisement call are homoplasious: the same pattern - *e.g*. a click-type call – recurs in genetically distant species.^6,27^ The temporal pattern of male advertisement calls is controlled by species-specific features of their hindbrain vocal pattern generating circuit.^28,29^ While temporal information can be ambiguous due to homoplasy, our data suggest that spectral information in advertisement calls – constrained by the morphology of the larynx – is more phylogenetically informative.^33^

In Xenopus, rapid oviposition once eggs are ovulated and the reproductive costs of hybridization provides a strong selective pressure for female recognition of same species’ male calls, especially when different species share the same pond. ^6^ The peripheral auditory system of females is tuned to their species’ own dyad: DF1, DF2 and the DF2/DF1 ratio.^34^ These features of vocal production and perception should reinforce the divergence of populations during speciation by limiting gene flow. *Xenopus* evolution however has also been shaped by multiple rounds of interspecific hybridization resulting in genomic introgression and the numerous highly polyploid species of the phylogeny, particularly A clade species.^4^ The acoustic advantage to a gravid female of locating the most genetically-compatible calling male using the clade-specific common harmonic dyad ratio could reflect co-evolution of the male vocal organ and auditory perception in the female.

## Methods

### Species

Subjects for this study were all sexually mature, male, clawed frogs from the sub-genus *Xenopus*. We either obtained frogs from commercial suppliers (Avifauna, Xenopus Express, Xenopus One, or Nasco) or from our *Xenopus* colony at Columbia University (species and populations: *X. pygmaeus*, *X. ruwensoriensis*, *X. amieti*, *X. boumbaensis*, *X. allofraseri*, *X. andrei*, *X. itombwensis*, *X. wittei*, *X. vestitus*, *X. lenduensis*, *X. largeni*, *X. gilli*, *X. poweri*, *X. laevis* Nigeria, *X. laevis* South Africa, *X. victorianus*, *X. petersii*, *X. laevis Malawi*, *X. borealis* and *X. muelleri*; see^6^ for details on geographic locales; species nomenclature as revised.^4^) Frogs were housed in 2-5 L of water in polycarbonate tanks, under a 12-12 light-dark cycle, fed frog brittle (Nasco; Ft. Atkinson, WI, USA) and had their water changed twice per week. All animals were housed and handled in accordance with the guidelines established by the Danish Animal Experiments Inspectorate (Copenhagen, Denmark) and the Columbia University Animal Care and Use Committee.

### Measuring arytenoid disc acceleration and velocity; sound pulse production

We used high speed films of tissue movements in the isolated larynx preparation^7^ of 5 adult male *X. laevis* to test the cavitation hypothesis. The animals were euthanized and their vocal organ and attached lungs were removed. The air-filled larynx was submerged in physiological saline in a 66 mm Petri dish. After isolation, larynges were pinned, ventral side up, via extra-laryngeal cartilages, to a Sylgard coated recording dish submerged in oxygenated Ringers solution. The bilateral, freed motor nerves were drawn into suction electrodes for stimulation (WPI Linear stimulus isolator model A395R-C). All isolated larynges produced sound pulses. Sound was recorded with a 1/2-inch pressure microphone-pre-amplifier assembly (model 46AD, G.R.A.S., Denmark) and amplified (model 12AQ, G.R.A.S., Denmark). The microphone and recording chain sensitivity was measured before each experiment (sound calibrator model 42AB, G.R.A.S., Denmark). The microphone was placed at 22-24 mm away from the mounted larynx. Because the signal to noise ratio of the hydrophone was lower than the microphone and the timing of sound events did not differ, we used the SNR microphone signal for further analysis. Microphone, hydrophone and stimulation timing signals were low-pass filtered at 10 kHz, (custom-built filter, ThorLabs, Germany) and digitized at 30 kHz (USB 6259, 16 bit, National Instruments, Austin, Texas).

Larynges were imaged with 12 bit high-speed camera (MotionPro-X4, 12 bit CMOS sensor, Integrated Design Tools, Inc.) mounted on a stereomicroscope (M165-FC, Leica Microsystems). The preparation was back-lighted to visualize the arytenoid discs by a plasma light source (HPLS200, Thorlabs, Germany) through liquid light guides and reflected of a 45° angled silver coated prism (MRA series, Thorlabs) to absorb heat. To track the position of landmarks, we placed 40-80 μm diameter carbon spheres on the surface of the larynx, muscles and arytenoid discs as illustrated in Fig. 1. The position of spheres was tracked at subpixel precision by interpolating the 2D intensity cross-correlations of the same sphere in an initial frame to each movie image (Fig 1B-D). Velocity and acceleration of the spheres were calculated by differentiation of their position. All control and analysis software was written in Matlab. In all 5 preparations, we filmed the larynx *in toto* following varying rates of nerve stimulation. In 3 preparations we obtained sufficiently high contrast images of the arytenoid discs at high imaging frame rates of 10,00-44,000 fps to allow automated position extraction during stimulation of the bilateral motor nerves at 40 Hz for 50 cycles.

Arytenoid gap width was defined as distance moved between the two markers from their resting position and perpendicular to the midline. Minimum and maximum acceleration of gap width were calculated per stimulus. Sound was bandpass filtered between 1-4 kHz (3th order butterworth filter implemented with zero phase-shift; filtfilt algorithm). The noise floor was defined as three times the standard deviation of a 67 ms background recording prior to each stimulation experiment. However, because sound energy did not fully dissipate in the experimental chamber between consecutive nerve stimulations, especially after 30-40 cycles, we used a threshold of 0.01 Pa to determine sound onset per stimulus. We used linear regressions - including only measurements above sound threshold – to calculate the minimal disc velocity and acceleration associated with sound generation.

### Recordings of vocal behavior

For *in vivo* recordings, frogs were placed in a 75 L aquarium. To promote vocal behavior, males were injected with human chorionic gonadotropin (hCG; Sigma: 50–200 IU depending on body size) one day prior to and on the day of recording. Males were then paired with a conspecific, sexually unreceptive female in a glass aquarium (60 × 15 × 30.5 cm, L×W×H; water depth = 23 cm; 20°C). A hydrophone (High Tech, Gulfport, MI, USA; output sensitivity -164.5 dB at 1 V/μPa, frequency sensitivity 0.015–10 kHz; or Cornell Bioacoustics, output sensitivity –163 dB at 1 V/μPa) was used to record calls to a Marantz digital recorder CD or flash card (CDR300, Marantz, Mahwah, NJ, USA; 44.1 kHz sampling rate) or on a computer (MacIntosh) via a Lexicon A/D converter. To measure values for DF1 and DF2, three non-consecutive advertisement calls (the smallest vocal unit as described in)^6^ were analyzed from 3 males of each species. Dominant frequencies were calculated from fast Fourier transforms (FFT) with maximum Q values (peak frequency/ maximum frequency 20 dB below peak frequency - minimum frequency 20 dB below peak frequency; Supplementary Table 1). The initial attack segment of each sound pulse was not included in the analysis because it is more broadband than the sustained portion of the call. The values shown in Supplementary Table 1 are the mean of individual means for all calls by species recorded.

### Sound and laser recordings in vivo and ex vivo

Advertisement calls of single males were recorded in aquaria with a hydrophone (H2a, Aquarian Audio Products; Anacortes, WA, USA) and vibrations were recorded simultaneously with a portable laser (PDV 100 laser, Polytec Inc.; Irvine, CA, USA) directed at the ventral surface of the singing frog. We recorded from one each of *X. laevis*. South Africa, *X. borealis*, *X. muelleri*, *X*. new tetraploid, and *X. boumbaensis*. We then isolated the larynx as described above. To access the elastic cartilages (Supplementary Figure SI1), we drilled a small hole in the dorsal surface of the larynx and then sealed it with a small piece of Parafilm. To ensure that the hole had no effect, sound and laser recordings were obtained before and after this procedure (Supplementary Figure SI1). The Parafilm was then removed and a 30g needle used to puncture the elastic cartilage on both sides (Supplementary Figure SI1), after which the Parafilm was replaced. At the end of the experiment, the larynx was split saggitally (‘butterflied’) and the disruption of elastic cartilage was confirmed by visual inspection (Supplementary SI1). Sound and laser recordings were digitized (Pre Sonus Audio box; Baton Rouge, LA, USA) and stored on a Macintosh computer.

## Acknowledgements

This work was supported by Danish Research Council (FNU) and Carlsberg Foundation to CPHE, a National Institutes of Health grant (R01 NS23684) to DBK, post-doctoral fellowships from the NIH (F32 GM103266) and the Revson Foundation to ICH and research funds associated with the Weintraub Chair (DBK). We thank Carolyn Diaz for laryngeal histology, Charlotte Barkan and Erik Zornik for reviewing the manuscript, and Sheila Patek and Ron Hoy for advice.

## Author Contributions

CPHE and DBK carried out the high-speed film recordings with the assistance of ICH. UKB heard the harmonic dyads. DOE, MLT and UKB obtained the *in vivo* and *ex vivo* sound and laser recordings. All authors contributed to designing experiments and analyzing the results. DBK and CPHE wrote the manuscript. All authors read and approved the final version of the manuscript.

